# Oncogenic NOVA1 expression dysregulates alternative splicing in breast cancer

**DOI:** 10.1101/2024.07.08.602566

**Authors:** Daniel F Moakley, Chaolin Zhang

## Abstract

Neuro-Oncological Ventral Antigen 1 (*NOVA1*) is best known for its role in mediating an alternative splicing (AS) program in neurons, yet was first discovered as an antigen expressed in breast tumors, causing rare autoimmune reactions and paraneoplastic neurological disorders (PNDs). The PND model suggests a plausible role of the tumor antigen expression in tumor suppression, whereas it has emerged that NOVA may function as an oncogene in a variety of cancers. In addition, whether NOVA mediates AS in breast cancer remains unanswered. Here we examine the AS profiles of breast invasive carcinoma (BRCA) tumor samples and demonstrate that ectopic *NOVA1* expression led to the activation of neuron-like splicing patterns in many genes, including exons targeted by NOVA in the brain. The splicing dysregulation is especially prevalent in cell periphery and cytoskeleton genes related to cell-cell communication, actin-based movement, and neuronal functions. We find that NOVA1-mediated AS is most prominent in Luminal A tumors and high *NOVA1* expression in this subtype is associated with poorer prognosis. Our results suggest that ectopic NOVA1 in tumors has regulatory activity affecting pathways with high relevance to tumor progression and that this might be a more general mechanism for PND antigens.

## Introduction

Paraneoplastic neurological disorders (PNDs) are rare autoimmune conditions in which patients exhibit neurological symptoms in conjunction with a cancer diagnosis. Tumoral expression of normally immune-privileged, neuron-specific onconeural antigens (ONAs) triggers an adaptive immune response targeting both the neoplasms and, in cases of PNDs, the patient’s central nervous system^1-8^. While ONA expression confers an immunogenic risk that causes neuronal degeneration, it was reported that the response can effectively curb cancer progression^9-13^. ONAs detected so far encompass a range of molecular and cellular functions^14-16^.Interestingly, several most common ONAs are RNA-binding proteins (RBPs) involved in regulation of neuron-specific RNA processing such as alternative splicing (AS)^5,6,15,16^, raising the possibility that they mediate aberrant neuron-like AS in tumors. However, such dysregulated AS has not been documented to the best of our knowledge.

Neuro-Oncological Ventral Antigen (NOVA) proteins were originally identified as the ONA recognized by the anti-Ri antibody found in serum or cerebrospinal fluid (CSF) of patients with paraneoplastic opsoclonus myoclonus ataxia (POMA) associated with breast, lung, or ovarian tumors, and characterized later to be neuron-specific RBP splicing factors^*1,2,6,17-19*^. The NOVA family consists of NOVA1 and NOVA2, which show partially reciprocal expression patterns in the central nervous system (CNS), with NOVA1 expression enriched in the midbrain, hindbrain, and spinal cord, and NOVA2 broadly expressed in the CNS but enriched in the neocortex and several other regions^*6,20*^. NOVA regulates a critical AS program in neurons^*21-24*^, with functional implications in neuronal migration^*25,26*^, axon outgrowth and guidance^*27*^, and synaptic formation^*25,28*^. Intriguingly, recent evidence has implicated NOVA1 as a potential oncogene affecting tumor proliferation, migration, tissue invasion, metastasis, and patient survival in several cancer types including hepatocellular carcinoma, melanoma, lung cancer, and glioblastoma^29-32^, although the opposite has also been suggested^33^. These studies in general point to NOVA1’s oncogenic roles enacted via gene regulation, including through dysregulated AS and inhibition of microRNA activities, as demonstrated in cell-based models^*30,32,34,35*^. However, whether tumoral *NOVA1* expression has AS regulatory activity in native tumors and how such activity is associated with tumor development and prognosis have not been assessed. In this report, we provide evidence that tumoral *NOVA1* expression mediates AS to facilitate a neuron-like splicing program correlated with poorer prognosis in a subset of breast invasive carcinoma (BRCA) neoplasms.

## Results

### NOVA expression across cancer types

To address the question of whether NOVA mediates AS in cancer, we first surveyed *NOVA1* and *NOVA2* expression using deep RNA-sequencing (RNA-seq) data from The Cancer Genome Atlas (TCGA), totaling 9,736 tumor and 726 normal samples across 33 cancer types (see Methods). We found *NOVA1* exhibited “runaway” expression in the majority of glioblastoma multiforme (GBM) and lower grade glioma (LGG), which have the brain origin, as well as a subset of breast invasive carcinoma (BRCA) samples, far exceeding that of the corresponding normal tissues (Supplementary Fig. 1a). In contrast, *NOVA2* was generally expressed at low levels and was rarely overexpressed in tumors (Supplementary Fig. 1b). This expression pattern, also noted in a recent study^32^ is consistent with the restricted expression of NOVA in the brain and also its known association with POMA in patients with breast cancer^2,15,36^.

### NOVA1 expression is associated with a neuron-like AS program in breast cancer

To assess NOVA1 splicing-regulatory activity in BRCA, we focused on cassette exons and performed differential splicing analysis using ten samples with the highest and lowest *NOVA1* expression levels, respectively (Fig. 1a). We detect 817 differentially spliced exons (DSEs) including 537 upregulated and 280 downregulated exons in *NOVA1*-high versus *NOVA1-*low samples using stringent thresholds (changes in percent spliced in (PSI), or |ΔPSI|≥0.2, false discovery rate (FDR)<0.05; Fig. 1b and Supplementary Table 1).

**Figure 1.**
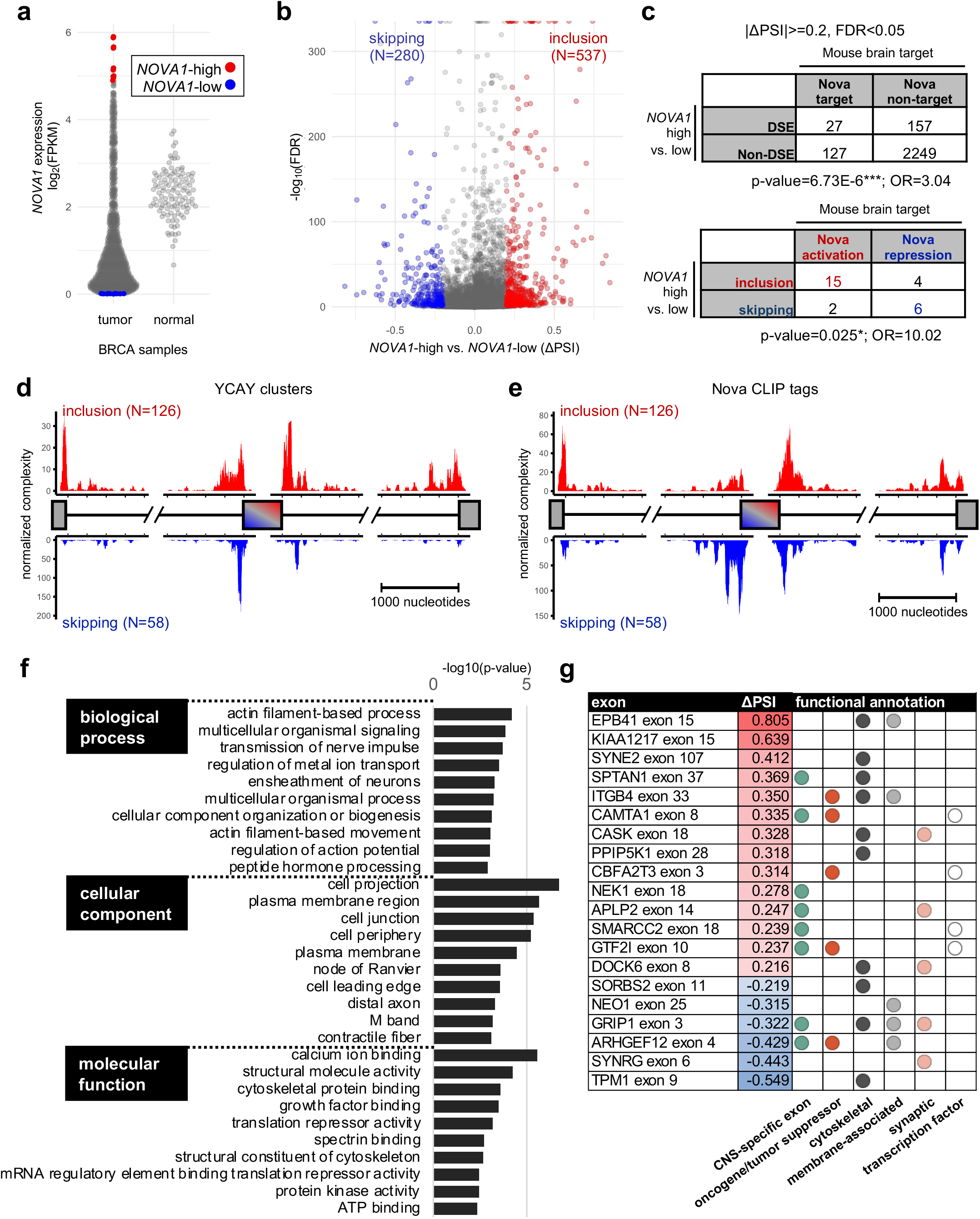
NOVA-mediated alternative splicing in breast cancer modulates CNS-specific, cytoskeletal, and surface membrane proteins associated with cell morphogenesis and motility. **a**, *NOVA1* expression of tumor and normal BRCA samples. *NOVA1*-high (N=10) and *NOVA1*-low (N=10) tumor samples selected for differential splicing analysis are indicated. **b**, Volcano plot of differential splicing analysis of *NOVA1*-high versus *NOVA1*-low BRCA samples. 537 upregulated and 280 downregulated differentially spliced exons (DSEs) were identified (|ΔPSI|>=0.2, FDR<0.05). **c**, Contingency tables showing significant overlap (top) and concordance of splicing-regulatory direction (bottom) of *NOVA1*-high versus *NOVA1*-low DSE mouse orthologs as compared to NOVA target exons in the mouse brain. **d**, RNA-map analysis showing the distribution of YCAY clusters surrounding mouse orthologs of upregulated (top) and downregulated (bottom) *NOVA1*-high versus *NOVA1*-low DSEs. **e**, Similar to (d) but for Nova CLIP tags. **f**, Top ten biological process, cellular component, and molecular function terms from GO analysis of genes containing of *NOVA1*-high versus *NOVA1*-low DSEs. **g**, CNS specificity, function annotations, and pathway associations of 20 genes containing the 21 high-confidence DSEs identified in *NOVA1*-high versus *NOVA1*-low BRCA samples with consistent NOVA-dependent splicing in the mouse brain. CNS-specific exons were determined by comparing brain samples to all other samples using GTEx RNA-seq data (see Methods).

To evaluate whether the DSEs are *bona fide* NOVA targets, we identified their orthologs in mice and compared them to a list of high-confidence NOVA target exons in the mouse brain that were previously determined by integrative modeling of NOVA-dependent splicing and direct NOVA-RNA interactions^23^. This analysis leveraged the observation that NOVA tends to regulate conserved AS events and many of the genes harboring neuron-specific exons have broad expression including BRCA. DSEs associated with NOVA1 expression overlapped significantly with the known NOVA mouse brain targets (Fig. 1c, top panel). Importantly, the direction of NOVA-dependent splicing changes observed in cancer samples is highly consistent with the direction of NOVA-dependent splicing regulation in the mouse brain (Fig. 1c, bottom panel). This overlap is robust when the a less stringent splicing change threshold (|ΔPSI|≤0.1) was used to include more NOVA target exons in cancer samples (Supplementary Fig. 2).

NOVA exhibits a stereotypical position-dependent binding pattern around its target exons (a.k.a. RNA-map), promoting exon inclusion when bound downstream of target exons and promoting skipping when bound upstream^21-23^. We examined evidence of NOVA binding around upregulated and downregulated DSE mouse orthologs and found an enrichment of both NOVA binding motif (YCAY, Y=C/U) clusters (Fig. 1d) and NOVA CLIP tags (Fig. 1e) in the expected locations, suggesting binding position-dependent splicing regulation. In addition, YCAY clusters predicted in the human genome also showed the expected pattern in BRCA *NOVA1-* high vs. -low DSEs without mouse orthologs, indicating that the tumors employ a partially human-specific NOVA-dependent splicing program (Supplementary Fig. 3). Together, these results demonstrate that tumoral expression of *NOVA1* elicited AS regulatory activity through direct protein-RNA interactions reminiscent of its activity in neurons.

### NOVA1-regulated DSEs in BRCA reside in CNS-specific, cytoskeletal, and cell surface proteins involved in cell migration

We next considered the possible biological functions of the dysregulated AS events observed in BRCA samples. DSEs significantly overlapped with CNS-specific alternative exons, consistent with the role of NOVA in establishing neuron-specific AS programs (Supplementary Fig. 4). Gene ontology (GO) analysis of the genes containing DSEs showed an enrichment for GO terms related to the cytoskeleton, cell-cell adhesion, cell-cell signaling, and neuronal functions including ensheathment, ionotropic signaling, and action potential regulation (Fig. 1f). Among the 21 DSEs that passed a strict inclusion change threshold (|ΔPSI|≥0.2) and matched the NOVA-dependent splicing change directions in the mouse brain, the majority show CNS-specific splicing and reside in a large proportion of cytoskeletal, membrane-bound, and synaptic genes. These include protein 4.1 (*EPB41*), a critical component of the membrane cytoskeleton that modulates physical properties of the membrane^37^, and integrin beta 4 (*ITGB4*), a known oncogene that lies on the boundary of the cytoskeleton and the extracellular matrix, modulates cancer cell organotropism, and correlates with tumor size and grade in BRCA^38,39^ (Fig. 1g). Also among them is Leukemia-Associated Rho Guanine Nucleotide Exchange Factor (*ARHGEF12)*, a Rho GTPase associated with prostate, lung, and breast cancers^40,41^ (Fig. 1g). Others are oncogenes, tumor suppressor genes, and transcription factors, suggesting modulation of different pathways classically associated with cancer (Fig. 1g).

### *NOVA1*-dependent AS dysregulation and decreased survival likelihood in Luminal A tumors

Gene expression profiling based on a panel of 50 cancer marker genes (termed “PAM50”^42,43^) has enabled BRCA tumors to be grouped into five subtypes: Normal-like, Basal-like, HER2-positive, Luminal A, and Luminal B. These approximately capture the clinical diagnostic groups that are differentiated based on histology, molecular profile, and the presence of estrogen (ER), progesterone (PR) or HER2 receptors, and correspond to different cellular compositions, mutational profiles, prognoses, and treatment options^44^. While roughly 10% of BRCA tumors molecularly resemble basal cells of the mammary ducts and are ER-, PR-, and HER2-negative, most express a luminal-like transcriptomic profile and at least one of the receptors^45,46^. Prognoses for luminal tumors are more favorable than basal tumors^45^, but they have heterogeneous types which can be more susceptible to recurrence^47^ and display various distinct metastatic profiles^45,48^.

To determine whether NOVA1 splicing activity is specific to particular tumor subtypes, we compared its expression among BRCA samples based on their PAM50 classification. High-*NOVA1* samples were enriched in Luminal A and Luminal B tumors, so we selected the highest and lowest expressing samples of those subtypes to repeat the splicing analysis (Supplementary Fig. 5a). High-versus low-*NOVA1* Luminal A and Luminal B samples showed splicing differences in 226 and 158 DSEs, respectively (|ΔPSI|≥0.2, FDR<0.05; Supplementary Table 1). Interestingly, orthologous DSEs in mice showed high concordance with the known NOVA targets for Luminal A but not Luminal B tumors (Fig. 2a and Supplementary Fig. 5b), suggesting that NOVA1-mediated AS dysregulation is most prominent in the Luminal A subtype.

We next asked whether high *NOVA1* expression was associated with prognosis in Luminal A and Luminal B BRCA. Survival analysis of patients with Luminal A tumors in the top 10% of *NOVA1*-expression showed significantly decreased survival compared to the other Luminal A cases (p=0.017 by log-rank test, Fig. 2b,c). The difference in survival is not significant for Luminal B tumors (p=0.36), although high *NOVA1* expression is also associated with a trend of reduction (Supplementary Fig. 5c). These data implicate *NOVA1* as an oncogene active in Luminal A tumors.

**Figure 2:**
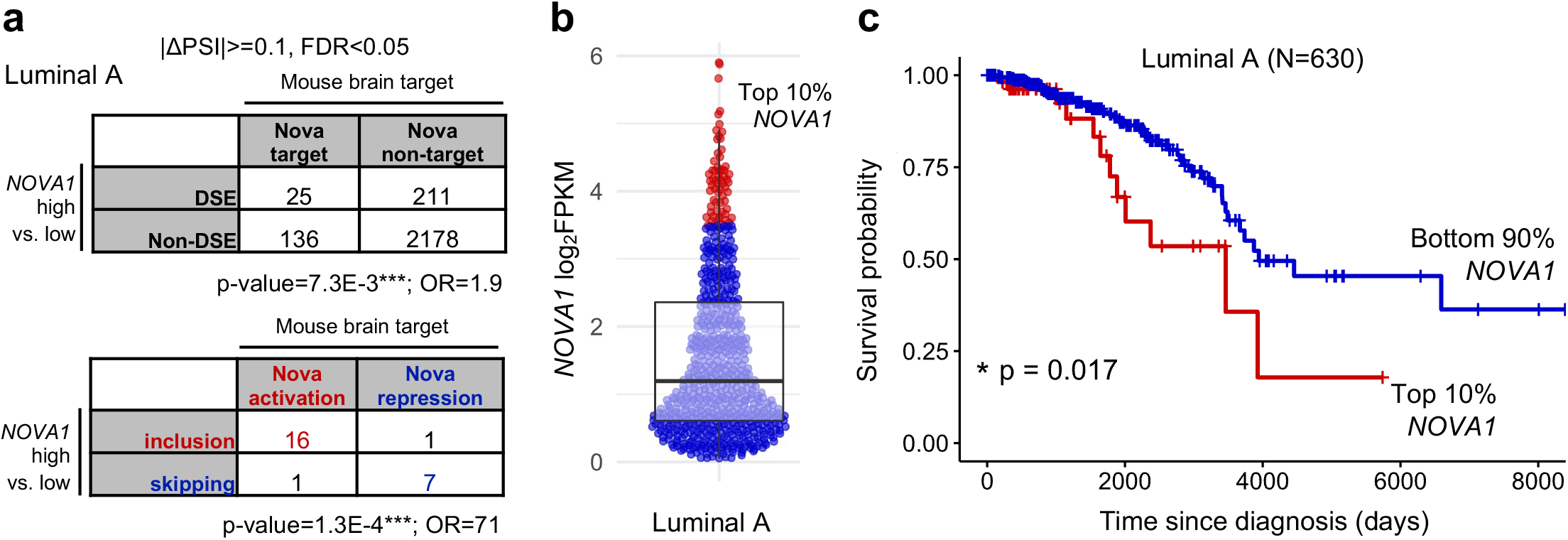
Evidence of NOVA-mediated AS dysregulation and prognosis in Luminal A breast tumors. **a**, Contingency tables showing significant overlap (top) and concordance in splicing-regulatory direction (bottom) of *NOVA1-*high versus –low Luminal A tumors. **b**, *NOVA1* expression in Luminal A tumors with the top 10%- and bottom 90%-expressing samples selected for survival analysis indicated in red and blue, respectively. **c**, Kaplan-Meier curve showing decreased survival in the top 10% versus bottom 90% *NOVA1*-expressing Luminal A tumors. P-value was calculated by log-rank test.

The observed NOVA-dependent AS dysregulation of genes involved in cell structure, communication, and motility raises the possibility that NOVA1 may enable tumor migratory and metastatic functions. The two main pathologic types of BRCA differ importantly in their tissue infiltration patterns: invasive lobular carcinomas (ILCs) which derive from mammary glandular cells, display a discohesive, single-file migration pattern that often escapes radiographic detection, whereas invasive ductal carcinomas (IDCs) form large, often calcified masses that are easily detectible on mammograms^49^. ILCs, which represent about ten percent of BRCA cases, mostly fall under the Luminal A subtype and express *NOVA1* significantly higher than IDCs (Supplementary Fig. 6a)^50,51^. *NOVA1* expression is also negatively correlated with that of E-cadherin (*CDH1*, Supplementary Fig. 6b), whose loss is a hallmark of ILC. Further work is warranted to determine whether NOVA1 is involved in ILC-specific pathology.

## Discussion

The PNDs are remarkable examples of naturally occurring tumor immunity in humans, as they provide an informative context to study effective tumor immunity, autoimmunity, and tumor biology. Such immunity was suggested to effectively suppress tumor growth so that the diagnosis of neurodegeneration frequently preceded cancer diagnosis. In small cell lung cancer, the detection of high-titer Hu antibody, which recognizes another neuronal RBP HuD/ELAVL4, was reported to be associated with better prognosis in multiple independent studies^9-12^. A conundrum arising the current PND model based on these observations is why tumor cells would ectopically express ONAs so that they can be recognized and attacked by the adaptive immune system. It was proposed that ONAs might be intrinsically oncogenic to provide selective advantage to the tumor cells^32^. This hypothesis is indeed supported by multiple reports suggesting NOVA acting as an oncogenic splicing factor when tested in cell-based models^30,32^. Our work extended these previous studies by providing evidence that tumoral expression of NOVA1 has splicing-regulatory activity in native breast tumors, which led to a large number of DSEs and the production of protein isoforms that are normally only expressed in the CNS. These genes are frequently involved in neuronal morphogenesis, cell-cell communication, and migration. These observations suggest a possibility that this dysregulated AS program may contribute to tumor migratory or metastatic functions, resulting in poorer prognosis compared to patients without NOVA1 expression in their tumors.

Why ectopic expression of ELAVL and NOVA proteins has apparently the opposite effects on tumor growth is unclear, but one possibility is worth considering. A moderate expression level of ONAs, which can be oncogenic and give selective advantage to tumor cells, might be insufficient to induce tumor immunity. However, a high level of ONA expression that mounts tumor immunity can cause tumor regression and give survival benefits to the patients (see model in Supplementary Fig. 7). We conjecture that ONAs might be oncogenic in general but different thresholds of ELAVL expression in small cell lung cancer and NOVA expression in breast cancer required to activate tumor-immunity might have contributed to the seemingly paradoxical observations on their contribution to tumor progression. When we extended our analysis to lung cancer, we found that in lung squamous cell carcinoma (LUSC), *ELAVL4* expression is associated with reduced survival (p=0.015; Supplementary Fig. 8), suggesting a potential oncogenic activity similar to NOVA1 in BRCA. Intriguingly, high doses of *ELAVL4* confer less risk than moderate doses (Supplementary Fig. 8b), perhaps indicating an activated tumor-suppressive immune response. Future analysis of the relationship between ONA expression, tumor immunity and outcome in small cell lung cancer (which is not included in TCGA) will help further to resolve the paradox.

## Author Contributions

Conceptualization: D.F.M., C.Z., data analysis: D.F.M., writing: D.F.M., C.Z.

## Methods

### RNA-seq data processing and analysis

RNA-seq data, gene expression tables, and accompanying tumor metadata were downloaded from the BRCA and LUSC projects in the TCGA database, generated by the TCGA Research Network (https://www.cancer.gov/tcga). Raw reads were mapped to the hg19 genome using OLego^52^. Quantification of TCGA-BRCA cassette exon inclusion and differential splicing test were performed using the Quantas pipeline (https://zhanglab.c2b2.columbia.edu/index.php/Quantas) as previously described^53^.

For Gene-Tissue Expression (GTEx) tissue cassette exon inclusion quantification, splicing junction read counts were downloaded from the GTEx consortium^54^. Splicing junctions were matched with the cassette exon annotations used in Quantas for splicing analysis to calculate PSI values. CNS-specific splicing was determined using ebayes^55^, a moderated t-test of mean PSI between brain areas and all other tissues.

### Nova-binding motif cluster, Nova CLIP tags, and Nova targets defined by integrative modeling

NOVA binding motif (YCAY) clusters used in Fig. 1d and Supplemental Fig. 3, NOVA CLIP data used in Fig. 1e, and NOVA mouse brain targets defined by the integrative Bayes network modeling used in Figs.1c, 2a, Supplementary Figs. 2 and 4 were obtained from our previous study^23^.

### Gene Ontology (GO) analysis

The R package “topGO”^56^ was used to perform Gene Ontology analysis. The “parent-child” algorithm was used for testing to control for term dependencies.

### Survival analysis

The R packages “survival”^57^ and “survminer”^58^ were used for survival analysis. Change in survival probability was assessed using the log rank test.

## Acknowledgments

We thank Zhang lab members and Robert B. Darnell for helpful discussion throughout the project. This study was supported in part by grants from the National Institutes of Health (NIH) (R01NS125018 and R35GM145279 to CZ). DFM was supported in part by NIH training grant T32NS064928.

## Supplementary Figure Legends

**Supplementary Figure 1. *NOVA1* and *NOVA2* expression in different types of tumors and control samples**. *NOVA1* **(a)** and *NOVA2* **(b)** expression in normal and tumor samples of various cancer types as quantified using TCGA RNA-seq data. Breast carcinoma (BRCA) samples are indicated with an arrow. Abbreviations for each cancer type are as follows: Adrenocortical carcinoma (ACC), Bladder Urothelial Carcinoma (BLCA), Breast invasive carcinoma (BRCA), Cervical squamous cell carcinoma and endocervical adenocarcinoma (CESC), Cholangiocarcinoma (CHOL), Colon adenocarcinoma (COAD), Lymphoid Neoplasm Diffuse Large B-cell Lymphoma (DLBC), Esophageal carcinoma (ESCA), Glioblastoma multiforme (GBM), Head and Neck squamous cell carcinoma (HNSC), Kidney Chromophobe (KICH), Kidney renal clear cell carcinoma (KIRC), Kidney renal papillary cell carcinoma (KIRP), Acute Myeloid Leukemia (LAML), Brain Lower Grade Glioma (LGG), Liver hepatocellular carcinoma (LIHC), Lung adenocarcinoma (LUAD), Lung squamous cell carcinoma (LUSC), Mesothelioma (MESO), Ovarian serous cystadenocarcinoma (OV), Pancreatic adenocarcinoma (PAAD), Pheochromocytoma and Paraganglioma (PCPG), Prostate adenocarcinoma (PRAD), Rectum adenocarcinoma (READ), Sarcoma (SARC), Skin Cutaneous Melanoma (SKCM), Stomach adenocarcinoma (STAD), Testicular Germ Cell Tumors (TGCT), Thyroid carcinoma (THCA), Thymoma (THYM), Uterine Corpus Endometrial Carcinoma (UCEC), Uterine Carcinosarcoma (UCS), Uveal Melanoma (UVM).

**Supplementary Figure 2. Concordance of *NOVA1-*associated BRCA DSEs with a more lenient inclusion change threshold** |Δ**PSI**|≥**0.1 with NOVA mouse brain targets**. Contingency tables showing significant overlap (top) and concordance in regulatory direction (bottom) of *NOVA1*-high versus *NOVA1*-low DSE orthologs and ground-truth Nova targets derived from Bayes integrative network modeling.

**Supplementary Figure 3. Evidence of *NOVA1*-mediated AS dysregulation for human-specific alternative exons**. RNA-maps show YCAY cluster distributions around all **(a)** or human-specific **(b)** *NOVA1*-high versus *NOVA1*-low DSEs.

**Supplementary Figure 4. BRCA *NOVA1*-high versus *NOVA1*-low DSEs are concordant with CNS-specific AS events**. Contingency tables showing significant overlap (top) and concordance in regulatory direction (bottom) of *NOVA1*-high versus *NOVA1*-low DSEs and CNS-specific exon inclusion or skipping (|ΔPSI|≥0.2, FDR<0.05).

**Supplementary Figure 5. *NOVA1*-high versus *NOVA1*-low Luminal B tumor DSE mouse orthologs are not concordant with Nova mouse brain targets. a**, Violin plots showing *NOVA1* expression in samples of the “PAM50” molecular tumor subtypes. For Luminal A and Luminal B subtypes, *NOVA1*-high (N=10) and *NOVA1*-low (N=10) tumor samples selected for differential splicing analysis are indicated in red and blue, respectively. The box plots indicate median and interquartile expression values. **b**, Contingency tables show overlap (top) and concordance in direction of regulation (bottom) of *NOVA1*-high versus *NOVA1*-low Luminal B DSE mouse orthologs and NOVA mouse brain targets. **c**, Kaplan-Meier curve showing decreased survival in the top 10% versus bottom 90% *NOVA1*-expressing Luminal B tumors. P-value was calculated by log-rank test.

**Supplementary Figure 6. Association of *NOVA1* expression with tumors of ILC pathology. a**, *NOVA1* expression in IDC versus ILC pathology types among BRCA tumors. The box plots indicate median and interquartile expression values. * p≤0.05, *** p≤ 0.001 (Student’s t-test). **b**, Scatter plot showing anticorrelation of *CDH1* and *NOVA1* expression among all BRCA samples. The Pearson’s correlation coefficient and the statistical significance are indicated.

**Supplementary Figure 7. A potential model of immune-tempered oncogenic function in ONA-expressing tumors**. While moderate ONA expression may confer oncogenic functions such as migratory and metastatic capabilities, high ONA expression can trigger an adaptive immune response that effectively inhibits tumor progression.

**Supplementary Figure 8. *ELAVL4* expression is associated with poor survival in an inverse dosage-dependent manner. a**, Violin plots showing *ELAVL4* expression in samples selected for survival analysis with groups indicated by color. Box plots indicate median and interquartile expression values. **b-d**, Kaplan-Meier curves showing decreased survival in *ELAVL4*-high vs. –moderate (b), *ELAVL4*-high vs. –negative (c), and *ELAVL4*-moderate vs. –negative (d) tumors. P-values are calculated by log-rank test.

