## Supplemental Figures S1-S8 for "Oncogenic NOVA1 expression dysregulates alternative splicing in breast cancer"

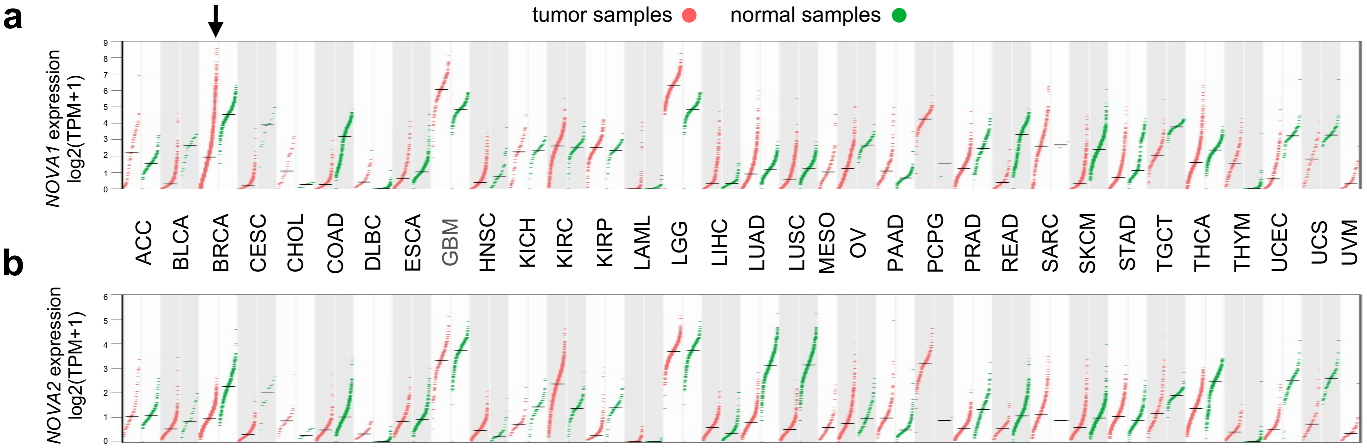
a
tumor samples
normal samples
NOVA1 expression log2(TPM+1)
ACC
BLCA
BRCA
CESC
CHOL
COAD
DLBC
ESCA
GBM
HNSC
KICH
KIRC
KIRP
LAML
LGG
LIHC
LUAD
LUSC
MESO
OV
PAAD
PCPG
PRAD
READ
SARC
SKCM
STAD
TGCT
THCA
THYM
UCEC
UCS
UVM
b
NOVA2 expression log2(TPM+1)

|ΔPSI|>=0.1, FDR<0.05

*NOVA1* high vs. low

| | Mouse brain target: Nova target | Mouse brain target: Nova non-target |
|---|---|---|
| **DSE** | 50 | 438 |
| **Non-DSE** | 104 | 1968 |

p-value=4.6E-5***; OR=2.2

*NOVA1* high vs. low

| | Mouse brain target: Nova activation | Mouse brain target: Nova repression |
|---|---|---|
| **inclusion** | 24 | 5 |
| **skipping** | 5 | 16 |

p-value=4.1E-5***; OR=14.3

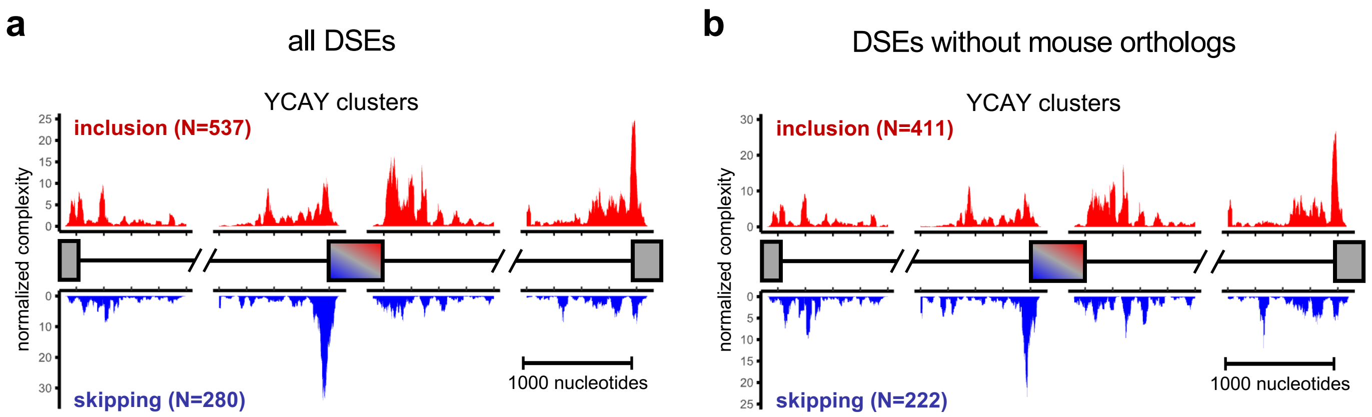
a
all DSEs
YCAY clusters
inclusion (N=537)
normalized complexity
25
20
15
10
5
0
0
10
20
30
skipping (N=280)
1000 nucleotides
b
DSEs without mouse orthologs
YCAY clusters
inclusion (N=411)
normalized complexity
30
20
10
0
0
5
10
15
20
25
skipping (N=222)
1000 nucleotides

|ΔPSI|>=0.2, FDR<0.05

| *NOVA1* high vs. low \ CNS vs. other tissues | DSE | Non-DSE |
|---|---|---|
| DSE | 156 | 661 |
| Non-DSE | 1689 | 14108 |

p=6.46E-12***

| *NOVA1* high vs. low \ CNS vs. other tissues | inclusion | skipping |
|---|---|---|
| inclusion | 67 | 34 |
| skipping | 22 | 33 |

p=0.0022***

**a**

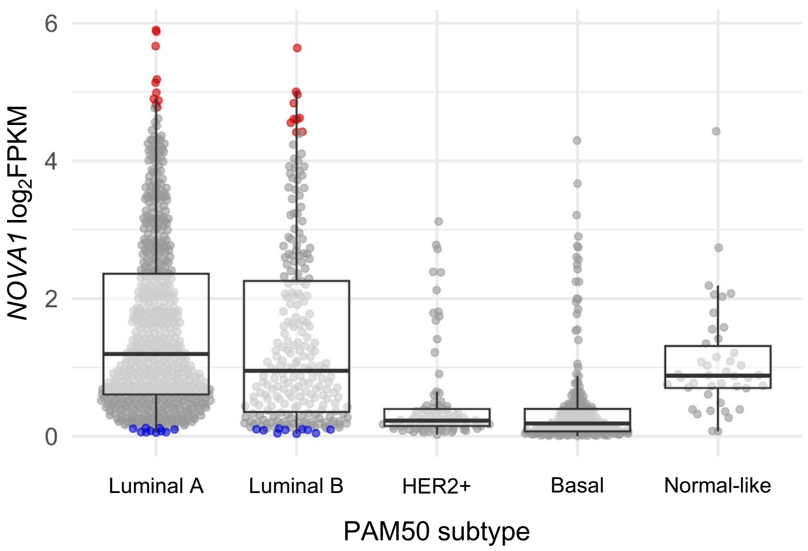


**b**

Luminal B

|ΔPSI|>=0.1, FDR<0.05

| NOVA1 high vs. low | Mouse brain target: Nova target | Mouse brain target: Nova non-target |
|---|---|---|
| DSE | 8 | 94 |
| Non-DSE | 142 | 2468 |

p-value=0.38; OR=1.39

| NOVA1 high vs. low | Mouse brain target: Nova activation | Mouse brain target: Nova repression |
|---|---|---|
| inclusion | 3 | 0 |
| skipping | 4 | 1 |

**C**

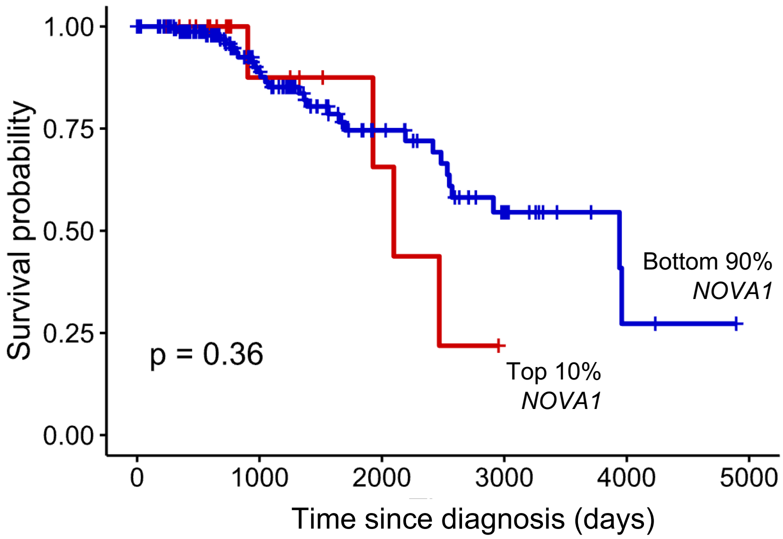

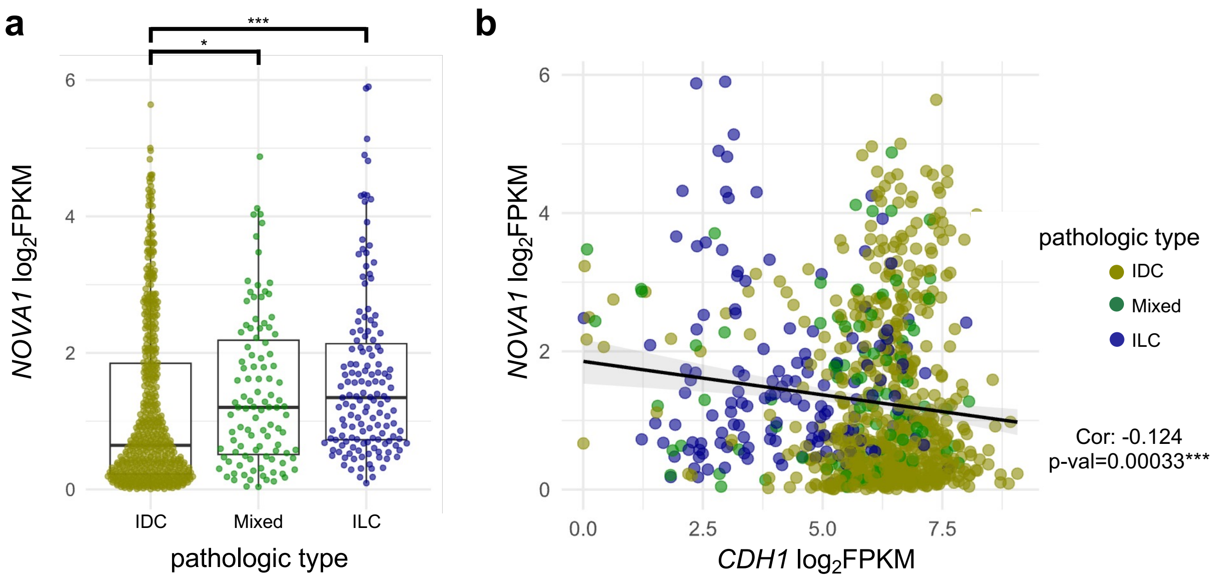

a
b
***
*
6
4
2
0
NOVA1 log2FPKM
IDC
Mixed
ILC
pathologic type
0.0
2.5
5.0
7.5
CDH1 log2FPKM
pathologic type
IDC
Mixed
ILC
Cor: -0.124
p-val=0.00033***

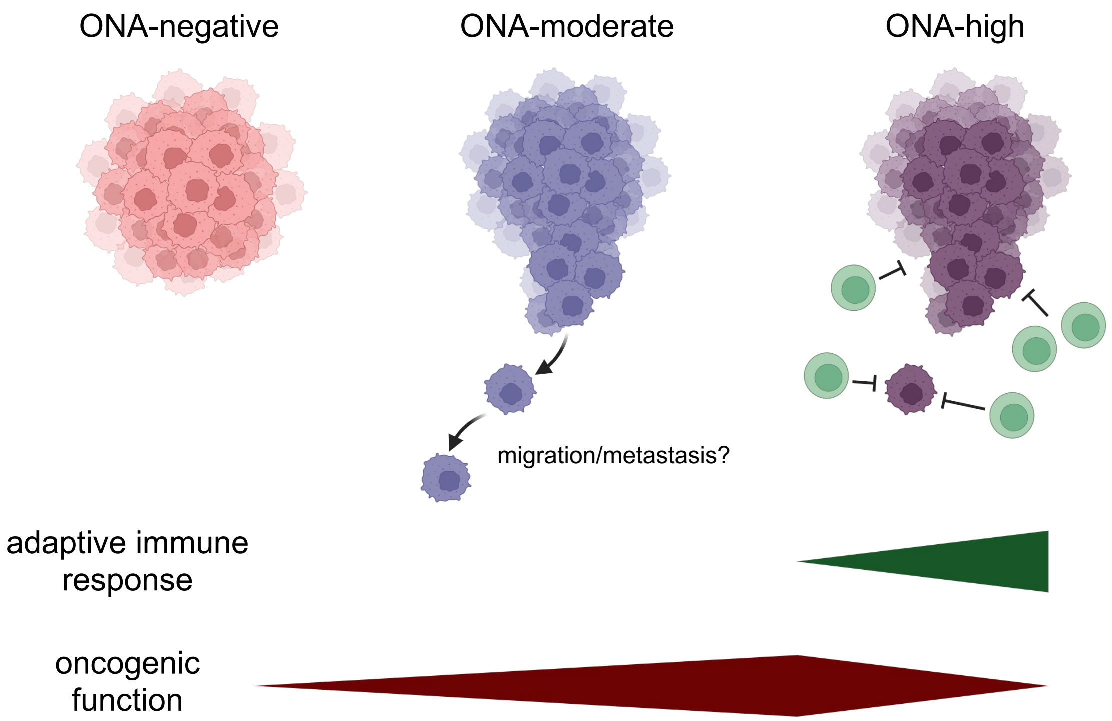
ONA-negative
ONA-moderate
ONA-high
migration/metastasis?
adaptive immune response
oncogenic function

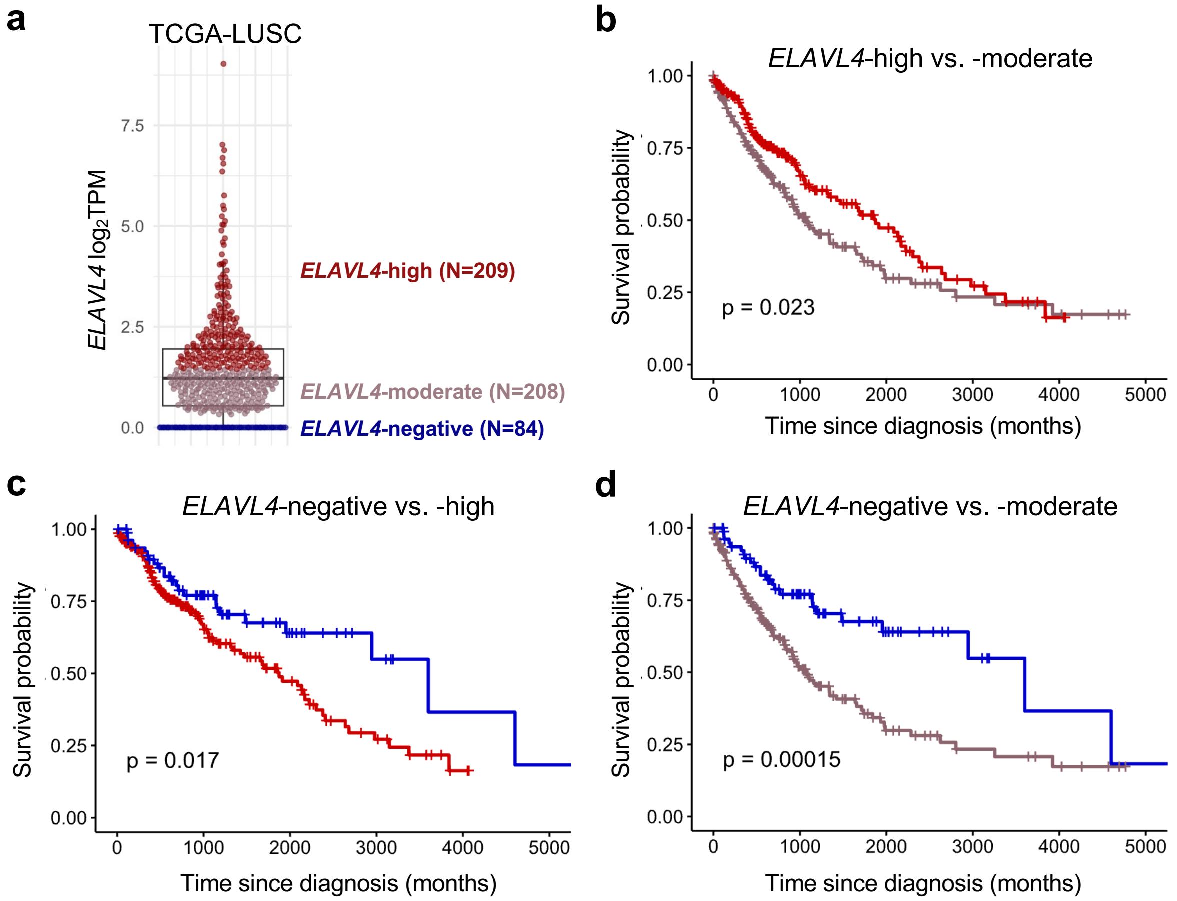
a
TCGA-LUSC
ELAVL4 log2TPM
7.5
5.0
2.5
0.0
ELAVL4-high (N=209)
ELAVL4-moderate (N=208)
ELAVL4-negative (N=84)
b
ELAVL4-high vs. -moderate
Survival probability
1.00
0.75
0.50
0.25
0.00
p = 0.023
0
1000
2000
3000
4000
5000
Time since diagnosis (months)
c
ELAVL4-negative vs. -high
Survival probability
p = 0.017
Time since diagnosis (months)
d
ELAVL4-negative vs. -moderate
Survival probability
p = 0.00015
Time since diagnosis (months)
